# Testing *in vitro* toxicity of nanoparticles in 3D cell culture with various extracellular matrix scaffold

**DOI:** 10.1101/2021.03.18.436024

**Authors:** Jae Won Choi, Song-Hwa Bae, In Young Kim, Minjeong Kwak, Tae Geol Lee, Min Beom Heo

## Abstract

Nanomaterials are used in a variety of fields and toxicity assessment is paramount for their development and application. Although most toxicity assessments have been performed in 2D (2-Dimensional) cell culture, the inability to adequately replicate the *in vivo* environment and toxicity is a limitation. To overcome the limitation, a 3D (3-Dimensional) cell culture method has been developed to make an environment closer to an *in vivo* system. In this study, 20 nm SiO_2_ nanoparticles were dispersed in serum-containing (SC) and serum-free (SF) media to compare 2D cell culture and 3D cell culture toxicity. The cells were subjected to a 3D cell culture method in which HepG2, a human-derived liver cancer cell line, was mixed on a scaffold. We found that nanoparticles induced toxicity in 2D cell culture, but toxicity was not observed in 3D cell culture similar to *in vivo* environment. However, differences in toxicity were observed between the three types of scaffolds in the absence of serum as the number of cells decreased.

## Introduction

Nanomaterials can be applied to diverse fields in a novel way or to provide enhanced functionalities depending on their particle size and surface modifications and therefore have been the subject of intense research in a wide range of fields, such as medicine, chemistry, biotechnology, foods, and electronics. As it has become possible to apply nanomaterials to various products and fields, many studies are underway investigating the effects that nanoparticle exposure may have on the body. In particular, silica nanoparticles are incorporated into various products, including drugs, cosmetics, food additives, and coatings, and are extensively studied in the biochemical field regarding their application as drug carriers and/or biomarkers [1, 2]. Cell cultures are used for such *in vitro* biochemical evaluation, and a variety of cell culture models and platforms have been developed to study investigational products in an environment that more closely mimic an *in vivo* system [3, 4]. Cell cultures are used in a wide range of fields to study various biochemical and physiological events that may occur *in vivo*, including the effects of drugs or toxic compounds on cells and development of mutations. Two-dimensional (2D) cell culture, which has been used since the early 1900s, is a cell culture method where cells are grown on flat surfaces optimized for cell attachment and growth. Traditionally, 2D cell cultures have been widely used, and they are still used in many studies because they allow simple cell observation and analysis. However, in 2D cell cultures, cells adhere and grow on the surface of a culture dish, and these cells assume a different morphology from cells grown *in vivo*. In addition, the 2D cultured cells differ from cells *in vivo* in a number of ways that may affect the *in vitro* tests, such as surface area exposed to the investigational product or cellular interactions [5]. S1 Fig shows the differences between 2D and 3D cell cultures. As a result, 2D cell culture methods have the limitation that they do not accurately reflect the *in vivo* environment, and as the cells grow and proliferate, the unnatural 2D environment can also affect gene expression [6-10]. To overcome the limitations of these 2D cell culture methods, a three-dimensional (3D) cell culture method has recently been developed and used to grow cells in 3D and create an environment similar to the *in vivo* conditions [11, 12]. To cultivate cells in 3D in vitro, cells must self-assemble to form cell aggregates, or scaffolds can be used to support 3D cell growth. Scaffold-based 3D cell culture technology typically uses a matrix system called extracellular matrix (ECM). Used for 3D cell culture, the ECM enables cell culture in a variety of spaces and can mimic cell-to-cell and cell-to-ECM interactions, including paracrine signaling [13]. ECM is composed of a variety of polymers such as collagen, enzymes, and glycoproteins and is known to support 3D cell growth and act as mediator for cell growth, migration, differentiation, survival, homeostasis, and morphogenesis [14-16]. In addition, moving away from 2D cell culture technology that uses culture dishes for cell culture, allows cells cultured in 3D, using various newly developed platforms, to be effectively applied to research. In this study, we compared the toxicity of silica nanoparticles in two models, using a widely used 2D cell culture model and a newly developed 3D cell culture model. Fig 1 shows a micro-pillar/micro well platform for building 3D cell culture techniques. Here, cells were mixed with ECM, dispensed into micro fillers, and cultured in micro wells. Here, the ECM serves as a scaffold to support cell culture in 3D. In addition, we evaluated the presence or absence of serum toxicity of ECM type nanoparticles using ECM (alginate extracted from algae, Matrigel extracted from mouse tail, and collagen extracted and purified from animals), and liver cancer cell line HepG2. Further, nanoparticle toxicity assessments were performed on cultured cells (1 × 10^3^ cells, 5 × 10^3^ cells, and 1 × 10^4^ cells) to assess differences in toxicity by ECM type.

**Fig 1.**
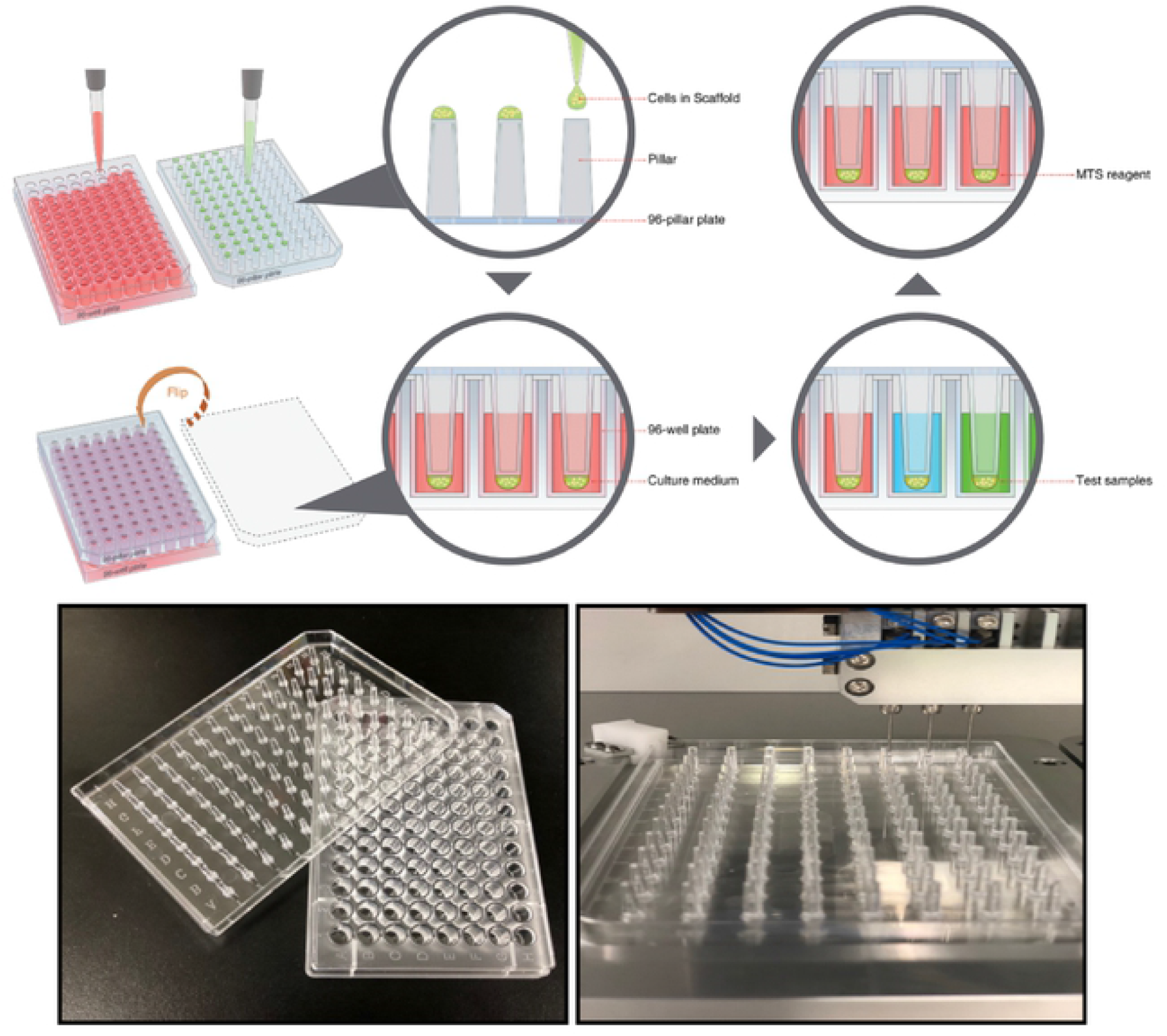
Micro Pillar / Micro Well Platform to Build 3D Cell Culture Technology.

## Materials and methods

### SiO_2_ nanoparticles

In the present study, 20 nm SiO_2_ nanomaterial, a certified reference material (CRM) developed by the Korea Research Institute of Standards and Science (KRISS), was used. The 20 nm SiO_2_ nanoparticles were prepared as previously reported [17]. The particles were measured using both Mobile Particle Size Meter (SMPS) and Dynamic Light Scattering (DLS). SMPS consisted of Differential Mobility Analyzer [(DMA), (TSI Inc., 3081)] and Condensation Particle Counter [(CPC), (TSI Inc.)] with a Brookhaven Instruments system having a BI9000AT digital correlator.

### Cell culture

The HepG2 human liver cancer cell line purchased from the American Type Culture Collection (ATCC) was used in this study. Prior to the experiment, HepG2 cells were thawed and allowed to acclimate for three cycles. In the culture medium, 500 mL of Dulbecco’s Modified Eagle’s Medium (DMEM) (Welgene™), 10% fetal bovine serum (HyClone™; FBS), and 1% penicillin-streptomycin (Welgene™; PS) were diluted to prepare a serum-containing medium (SC). Next, 15 mL of previously prepared DMEM was poured into a sterile T75 flask purchased from Corning^®^. Cells were harvested in T75 flask with 0.05% trypsin-EDTA (1X) purchased from Sigma Aldrich. Thereafter, cells were counted using Trypan blue 0.4% purchased from Sigma Aldrich and placed in T75 flasks containing 15 mL DMEM at 2 × 10^6^ cells/mL for incubation (37 °C/ 5% CO_2_/ 95% humidity) for 2 days to reach approximately 80–90% confluence. Serum-free media (SF) was prepared by adding only 1% penicillin-streptomycin (Welgene™; PS) to 500 mL of Dulbecco’s Modified Eagle’s Medium (DMEM) (Welgene™).

### 2D cell culture and 3D cell culture in scaffold

S2 Fig shows the complete protocol of 2D and 3D cell culture. First, the HepG2 cells were cultured in T75 flasks for 2 days, the culture medium was removed, and the cells were washed once with phosphate buffered saline (PBS), 3 mL of 0.05% trypsin-EDTA (1X) was added, and the cells were incubated for 2 min (37 °C/ 5% CO_2_/ 95% humidity). To each T75 flask containing cells, 7 mL of DMEM was added and the detached cells were centrifuged for 5 min at 2500 rpm. The supernatant was removed and the cell pellet was resuspended in 1 mL of DMEM. Thereafter, the cells were counted and seeded at a density of 1.0 × 10^4^ cells/well in 200 μL suspension in the wells B–G (3-6, 8-10) of a 96-well plate for 2D culture. S3 Fig (a) shows a layout protocol in which 2D and 3D cell cultures were made to form in the B–G (3-6, 8-10) column. We prepared three plates in which 2D HepG2 cells were cultured. Further, 200 μL of PBS was added to wells in A and H columns to prevent evaporation of the culture medium from the plate during culture. For 3D cell culture, cells were initially cultured in the same way as 2D cell culture. In 3D cell culture, a 96-pillar plate (micro-pillar/micro well) consisting of 0.2 mm diameter pillars was used, and cells were seeded in the same order as seeded for 2D cell culture. Equal number of cells (1.0 × 10^4^ cells/pillar/2 µL) was used to compare the toxicity levels in 3D and 2D cell cultures. To evaluate the toxicity in different types of scaffolds used in 3D cell culture, cells were grown at densities 5.0 × 10^3^ cells and 1.0 × 10^3^ also. In this study, three types of ECM (alginate, Matrigel, and collagen) were used as scaffolds for 3D cell growth. The first type of scaffold is 3% alginate, which is liquid at 4 °C, but forms a gel at room temperature. To dilute 3% alginate to a final concentration of 0.75%, 3% alginate was diluted 1:1 in DMEM and further diluted in 1:1 ratio in DMEM containing the prepared cells. To culture three plates for each cell density, an automated 3D cell culture system (MBD model) was used to place the cells on a 96-pillar plate and let the gelation proceed for 5 min at room temperature. Cells were placed in a 96-well plate containing 200 µl of DMEM prepared in advance and incubated for 24 h (37 °C/ 5% CO_2_/ 95% humidity). Similarly, the 3D cell culture was carried out using two other scaffolds: 100% Matrigel (Corning^®^) and collagen (Corning^®^). Matrigel is a liquid at 4 °C and quickly turns into a gel at room temperature. Therefore, it was placed on ice for use in experiments. Here, Matrigel was diluted in DMEM containing cells in a 1:1 ratio to grow cells in 3D at 50% Matrigel concentration. Equal number of cells was seeded in the same order as alginate; cells were transferred from the incubator to an empty 96 well plate, preheated, and gelation proceeded for 10 min. Then, cells were transferred to a 96-well plate containing 200 µL of DMEM, and cultured for 24 h (37 °C/ 5% CO_2_/ 95% humidity).

### Nanoparticle and chemical control process

A CRM developed by the Korea Research Institute of Standards and Science (KRISS), 20 nm SiO_2_ nanoparticles dispersed in sterile distilled water at a concentration of 10 mg/mL, was used. As described above, the effect of corona protein was taken into consideration, SC and serum-free (SF) media were separately prepared to have a final 20 nm SiO_2_ concentration of 1000 μg/mL. To compare the toxicity of nanoparticles, CdSO_4_ (Cadmium sulphate) in the powder form purchased from Sigma-Aldrich was used as the chemical control at varied concentrations of 0, 9.4, 18.8, 37.5, 75 and 150 μM. In addition, CRM 20 nm SiO_2_ nanoparticles at a concentration of 9.4 mg/mL were used and diluted to 0, 10, 50, 100, 500, 1000 μg/mL in 2D cell culture. In 3D cell culture, higher concentrations of 0, 62.5, 125, 250, 500, and 1000 μg/mL were prepared considering the penetration of nanoparticles into the scaffold. Prior to experimentation, 20 nm SiO_2_ diluted to each concentration in SC and SF media was vortexed for 30 s to evenly disperse the particles. Using an electronic scale, 103 mg of CdSO_4_ was added to 40 mL of distilled water and vortexed for 30 s to prepare 10 mM CdSO_4_, which was diluted to a final concentration of 150 μM and then further diluted in two-folds to prepare various concentrations mentioned above. In this study, cytotoxicity was confirmed using a toxicity test method based on absorbance measurement.

### Cell viability measurement through MTS assay

After exposure of cells to nanoparticles for a 24 hr of time, toxicity was quantified by MTS assay, which is one of the methods generally used to measure cell proliferation or toxicity [18]. CellTiter96® AQueous One Solution Cell Proliferation Assay kit purchased from Promega was used. MTS reagent was diluted in each culture medium (SC and SF) used in the experiment at a ratio of 1:5 (MTS reagent: media), and 120 µL of the mixture was added to the cells following incubation. In 3D cell culture, diluted MTS reagent was added directly to an empty 96-well plate and was transferred to the 96-pillar plate in which cells were seeded. Cells were incubated with MTS reagent for 1 h for the 2D cell culture group and 2 h for the 3D cell culture group considering the effects of ECM. The reaction was allowed to proceed in an incubator (37 °C / 5% CO_2_ / 95% humidity). The absorbance of these 96-well plates was measured at 490 nm using a micro-plate reader. Cell viability was calculated from the following equation, and IC_50_ values were calculated using SoftMax Pro software.

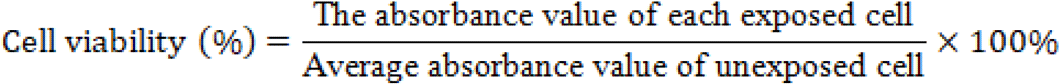

## Statistical Analysis

Results of all the experiments were statistically analyzed and represented as the mean with the standard error of the mean (SEM) of three or more independent experiments (n = 3). The t-test was performed by dividing the equal variance and this variance through the F-test using Excel, and the *p-value* was statistically processed as a value expressed in both directions. In the 2D cell culture statistical comparisons were performed between two groups, and in 3D cell culture, three groups were compared. While comparing the scaffolds, alginate was marked with * and Matrigel with #. All *p-values* are expressed * as *p*<0.05, ** as *p*<0.01, and *** as *p*<0.001. IC_50_ (inhibitory concentration 50) values were calculated using the SoftMax Pro program.

## Results

### Size changes in 20 nm SiO_2_ with or without serum

The particle size distribution is plotted in Fig 2 (a) and (b) showing hydrodynamic size and mobility diameter, respectively. Fig 2 (c) and (d) respectively, show transmission electron microscopy (TEM) images of 20 nm SiO_2_ acquired on JEOL JEM-ARM200F and scanning electron microscopy (SEM) images of 20 nm SiO_2_ acquired at 200 kV accelerating voltage, at an acceleration voltage of 10 kV each on the ZEISS Gemini SEM 500. As shown in Fig 2 (e), particle size measurement by various methods revealed that the particle size data for 20 nm SiO_2_ obtained by electron microscopy (TEM and SEM) were similar to that obtained by hydrodynamic methods (DLS and SMPS). This indicates that the nanoparticles were highly monodisperse in aqueous suspension without aggregation.

**Fig 2.**
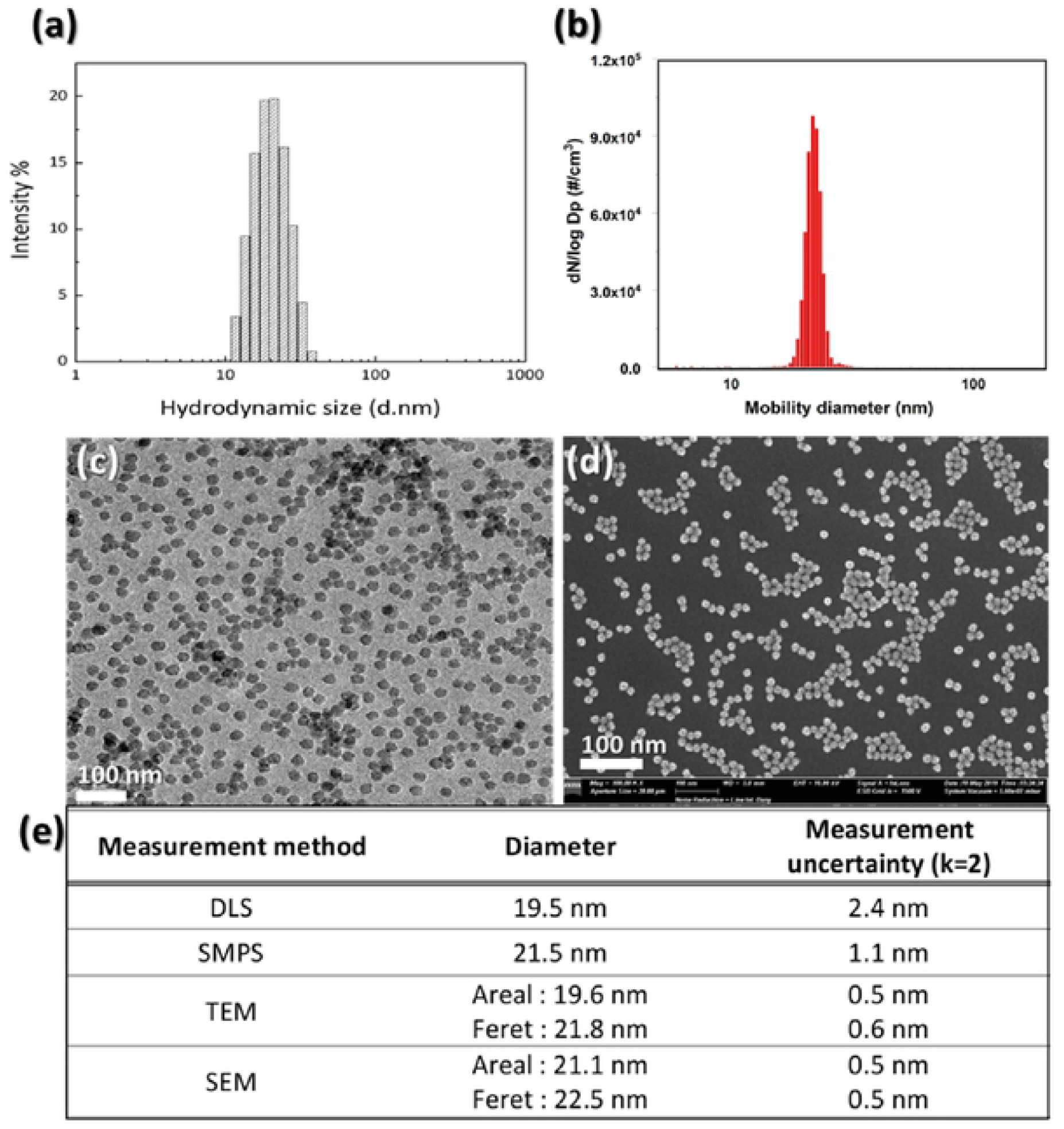
Size Analysis of SiO_2 N_anoparticles by Different Methods. **(a)** Graphical representation of Z-average diameter of 20 nm SiO_2_ (avg 19.5 ± 2.4 nm) analyzed using DLS. **(b)** Graphical representation of Z-average diameter of 20 nm SiO_2_ (avg 21.5 ± 1.1 nm) analyzed using SMPS. **c)** Representative TEM image captured at an acceleration voltage = 300 kV. **(d)** Representative image of 20 nm SiO_2_ nanoparticles observed using SEM. **(e)** Table shows the size of 20 nm SiO_2_ particles measured using different approaches.

To determine the cytotoxicity of the 20 nm SiO_2_ nanoparticles, they were mixed with culture media to expose cells to the nanoparticles. In addition, it has been reported that proteins or polymers present in culture media get adsorbed to the nanoparticles and form protein corona that can alter the size of the particles in an aqueous solution [19, 20]. Therefore, prior to experimentation, particle size in the culture media with or without serum, in which the cells were exposed to the nanoparticles, was determined using DLS. As shown in Fig 3, we were able to confirm that the size of 20 nm SiO_2_ nanoparticles dispersed in SC media increased to as high as 158 nm because the serum led to the formation of protein corona around the nanoparticles. In contrast, we found that the size of 20 nm SiO_2_ nanoparticles dispersed in SF media was 23 nm, which was similar to the original size.

**Fig 3.**
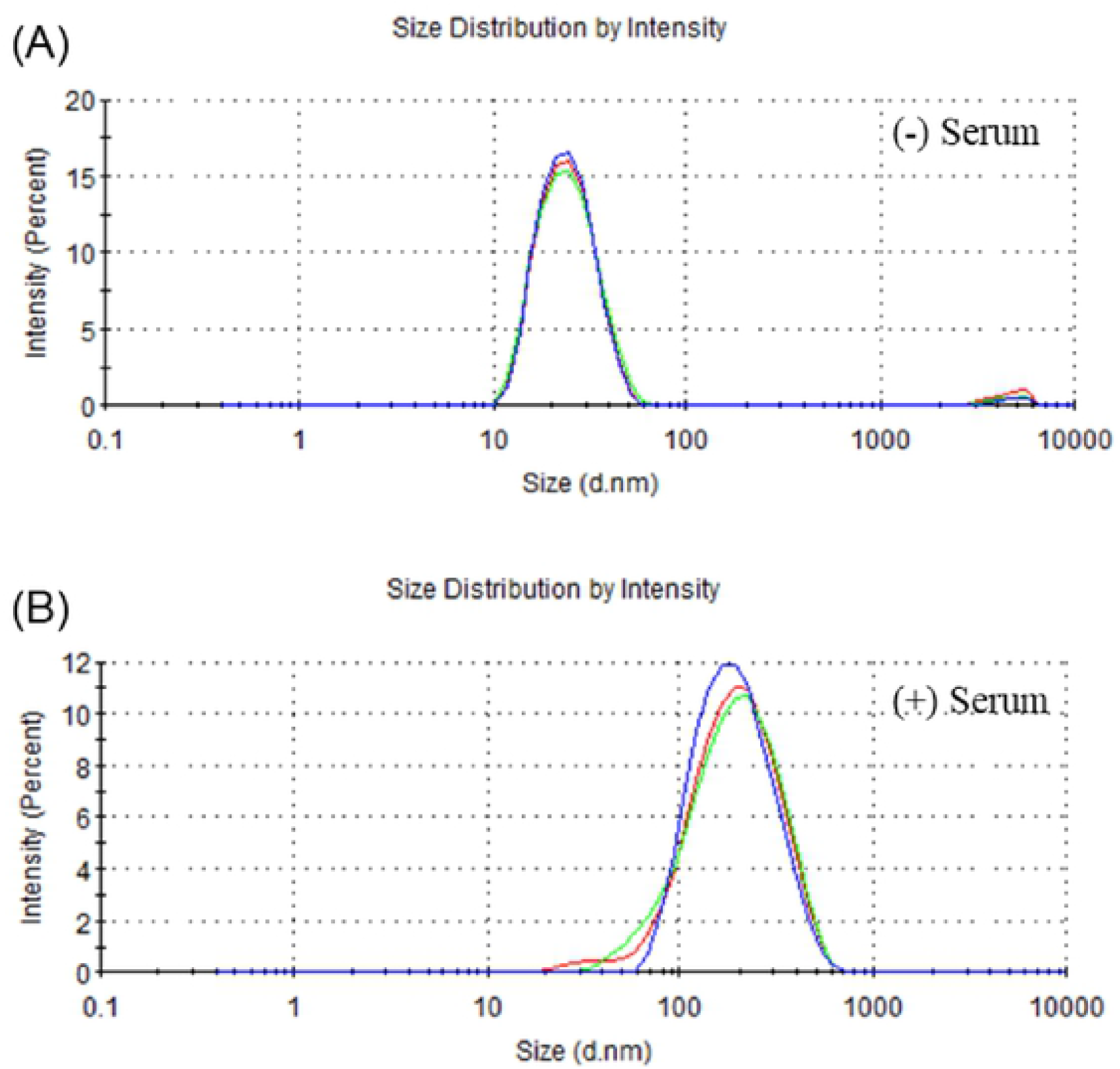
Size of 20 nm SiO_2_ Nanoparticles Dispersed in DMEM With and Without Serum Analyzed Using DLS. **(a)** The size in serum-free DMEM is 23.23 ± 0.39 nm **(b)** The size in serum-containing DMEM is 157.07 ± 1.07 nm.

### Comparison of cell growth differences according to types in ECM (Scaffold)

In this study, cell growth rates in three types of ECM-alginate, Matrigel, and collagen were compared prior to the nanoparticle toxicity evaluation test. For a more reliable comparison, cells were cultured in SF media all day in an incubator (37 °C/5% CO_2_/95% humidity) and then observed with MTS analysis and Optical microscopy. Fig 4 shows 3D cultured HepG2 cells on each scaffold observed after 24 h.

**Fig 4.**
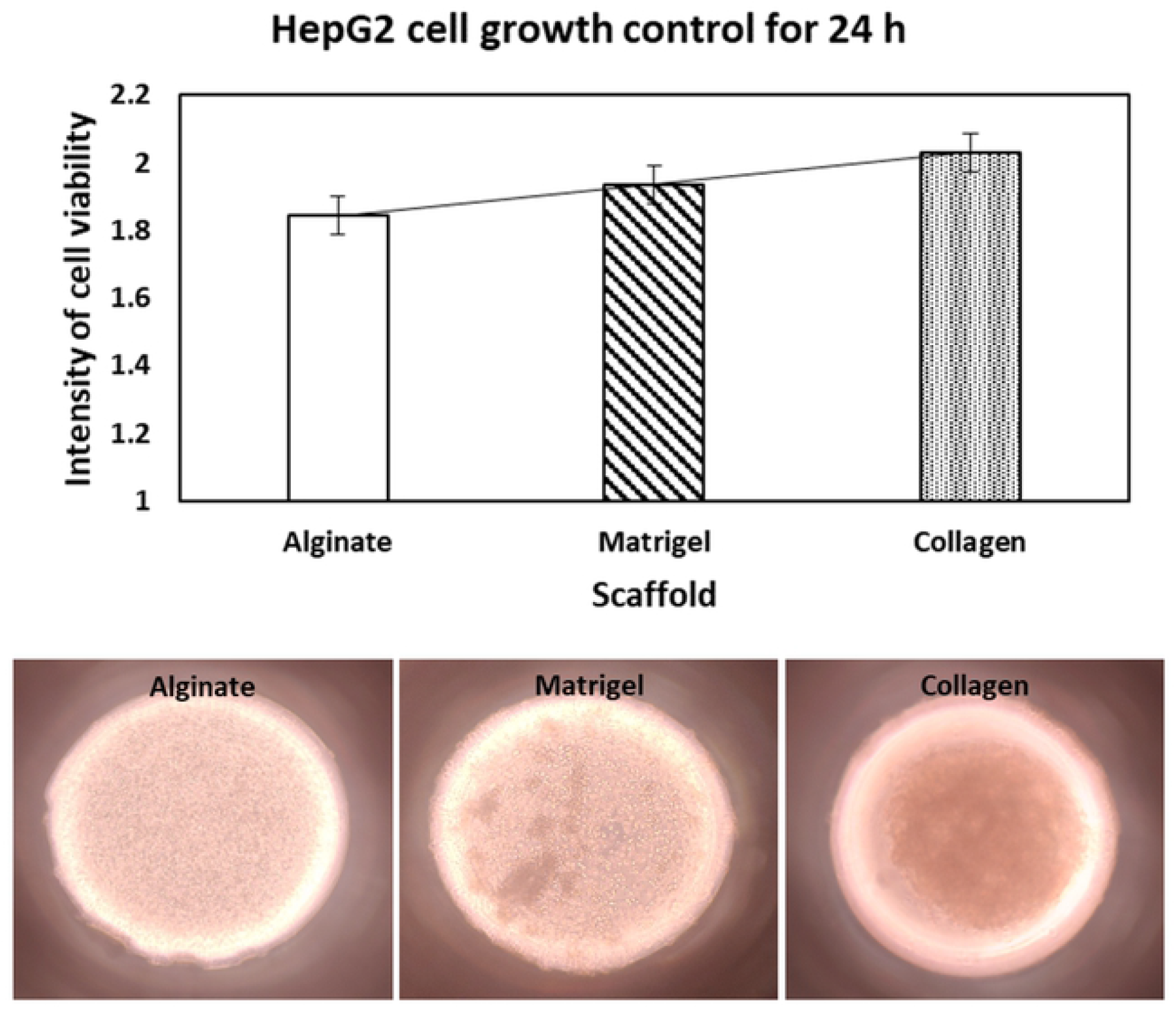
Different Types of Scaffolds (alginate, Matrigel, and collagen) Showed Differences in the Growth of HepG2 Cells Under Serum-free Media Conditions for 24 h.

Observation under Optical microscope showed that in the case of alginate, the cells form a single spheroid, and in the Matrigel, the cells gradually combine to form a small spheroid. However, collagen appears to constitute a spheroid in which cells are completely coagulated. MTS analysis showed that cell growth was promoted by collagen rather than alginate and Matrigel. These results were similar to those reported previously, stating that collagen promotes cell growth and is a useful ECM [21].

### Comparison of 2D and 3D cytotoxicity of nanoparticles with and without serum

The HepG2 cells grown in 2D and 3D modules were treated with varied concentrations of SiO_2_ nanoparticles (0 to 1000 µg/mL) and a chemical control-CdSO_4_ (0 to 150 µM) dispersed in SC and SF media. S3 Fig (b) shows nanoparticles and chemical control prepared for each concentration. The nanoparticles prepared for each concentration were added to 96 well plates (B–G, 8–12); among them, the wells without cells (B–G, 11–12) were used for nanoparticles interference control. CdSO4, a chemical control, was also treated in 96 well plates (B–G, 1–5), and treatment in wells without cell (B–G, 1–2) acted as the chemical interference control. In addition, only media replaced cell control in B–G, column 6 of the 96 well plate. No cells present in B–G, row 7 of a 96 well plate, acted as the background media control. MTS assay was used for toxicity analysis, and the results were quantified using a microplate reader. Based on the quantified data, IC_50_ (inhibitory concentration 50) values were calculated using the SoftMax Pro program to determine the concentration at which 50% cell death occurred. Fig 5 shows the cell viability of the 2D cultured HepG2 cells after exposure to CdSO_4_ and 20 nm SiO_2_ nanoparticles in media with or without serum.

**Fig 5.**
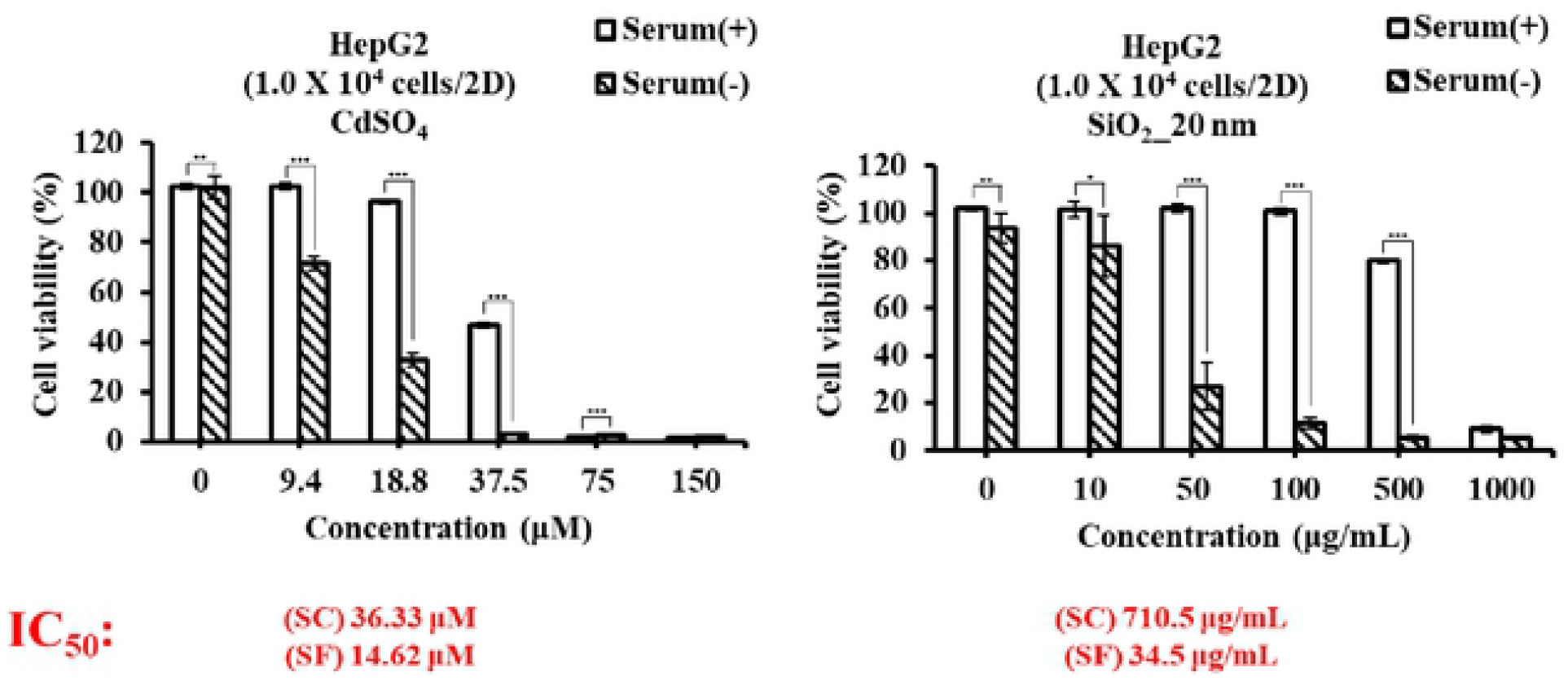
Toxicity Analysis of CdSO_4_ and 20 nm SiO_2_ Nanoparticles Dispersed in DMEM With or Without Serum Against 2D Cultured HepG2 Cells Analyzed by MTS Assay. The data is represented as standard error of the mean (SEM) of 3 independent experiments (n = 3) statistical significance is analyzed by t-test. The *p-value* is denoted as* *p <0*.*05*, ** *p <0*.*01*, and *** *p <0*.*001*.

For CdSO_4_, the IC_50_ value was 36.33 μM in the SC and 14.62 μM in SF media, indicating that in the absence of serum, IC_50_ value was lowered, indicating that it caused more toxicity. In the 2D cell culture model, the IC_50_ value of the toxicity of nanoparticles was 710.5 μg/mL in the SC, and 34.5 μg/mL in SF media. Similarly, toxicities were compared when three types of scaffolds (ECM) at 1.0 × 10^4^ cell density were used for HepG2 3D cell culture as shown in Figs 6 and 7. First, in the chemical control CdSO_4_, the IC_50_ in alginate was 70.3 μM in the SC and 29.17 μM in SF media. The IC_50_ value in Matrigel was 108.6 μM in the SC and 21.09 μM in SF conditions. In addition, the IC_50_ value in Collagen was 94.5 μM in the SC and 21.87 μM in SF media. For the toxicity of nanoparticles, the IC_50_ values of alginate, Matrigel, and collagen could not be determined regardless of the presence or absence of serum. These results suggest that in the presence of serum, protein coronas are formed around the nanoparticles, increasing the particle size, preventing the particles from entering the HepG2 cells in the scaffold, and thus reducing the toxicity compared to 2D cultured cells. This suggests that even in the absence of serum, cytotoxicity was markedly reduced in 3D cell culture due to the influence of cell resistance and scaffold. In addition, the results obtained from 3D cell cultures using synthetic ECM as scaffolds to create conditions similar to the *in vivo* environment are shown to differ from those obtained from 2D cell cultures traditionally used for *in vitro* studies.

**Fig 6.**
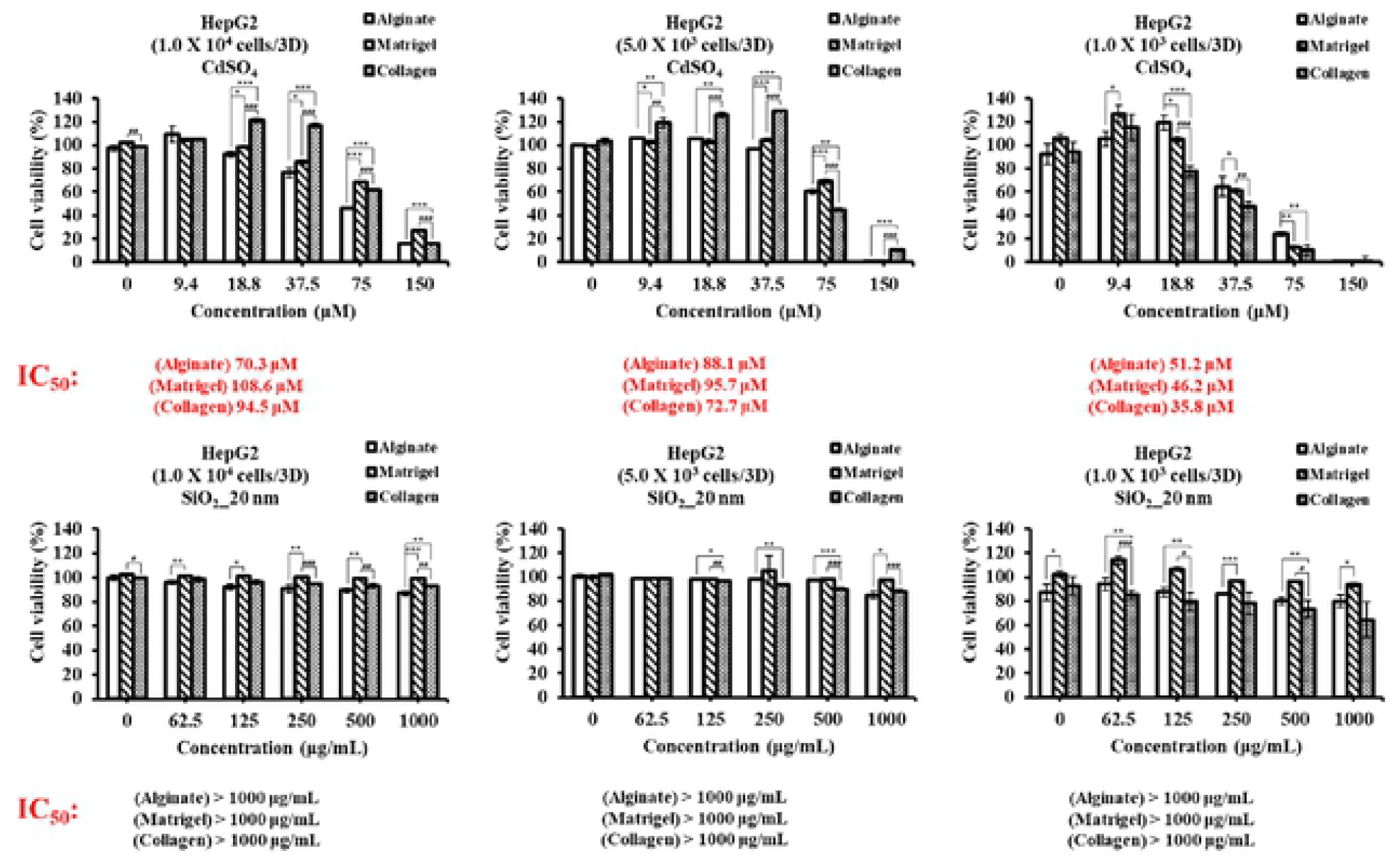
Toxicity of 20 nm SiO_2_ Nanoparticles Under Serum Containing Media Conditions Was Compared by Changing the HepG2 Cell Count for Each Type of Scaffold (alginate, Matrigel, and collagen). The standard error of the mean (SEM) of 3 independent experiments (n = 3) was statistically analyzed as a *p-value* via t-test. The *p-value* is denoted as* *p <0*.*05*, ** *p <0*.*01*, and *** *p <0*.*001*. Comparisons with alginate are indicated by *, and comparisons with Matrigel are indicated by #.

**Fig 7.**
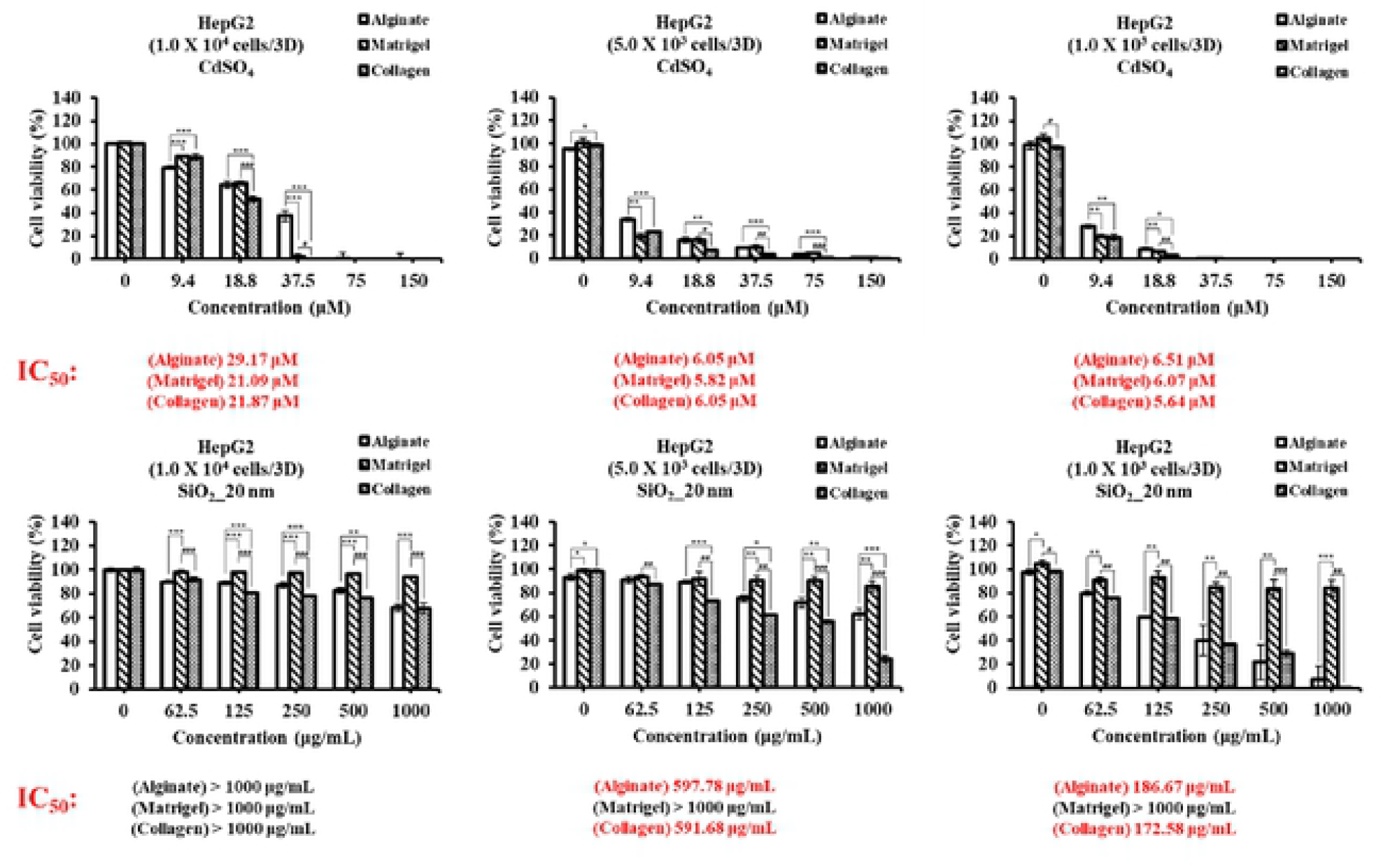
Toxicity of 20 nm SiO_2_ Nanoparticles Under Serum Free Media Conditions Was Compared by Changing the HepG2 Cell Count for Each Type of Scaffold (alginate, Matrigel, and collagen). The standard error of the mean (SEM) of 3 independent experiments (n = 3) was expressed as a *p-value* via T-test. The *p-value* is denoted as* *p <0*.*05*, ** *p <0*.*01*, and *** *p <0*.*001*. Comparisons with alginate are indicated by *, and comparisons with Matrigel are indicated by #.

### Comparison of cytotoxicity of nanoparticles in different 3D scaffolds

Since standards for 3D cell culture have not yet been established, various 3D cell culture methods have been adopted and various hydrogels have been developed for use as ECM. Therefore, we created conditions for growing HepG2 cells as spheroids in 3D cell culture using three ECM, alginate, Matrigel, and collagen, which are commonly used as scaffolds in 3D cell culture. The effect of ECM types on the cytotoxicity of nanoparticles was compared in the same way as in the above experiment using 3D cell culture plates grown on different ECM. In addition, the difference in toxicity of 20 nm SiO_2_ to 3D cell culture was confirmed by the presence or absence of serum. Fig 6 shows the toxicity of CdSO_4_ and 20 nm SiO_2_ nanoparticles as a function of the number of HepG2 cells grown on three different scaffolds in the presence of serum. In the presence of serum, the size of the nanoparticles is increased by a large number of proteins present in the medium getting adsorbed on the particles, increasing the original size of the nanoparticles. Therefore, cell uptake has not taken place, resulting in no toxicity. In contrast, for the control chemical CdSO_4_, IC_50_ values can be obtained on three different scaffolds as the concentration increases. In addition, it was confirmed that smaller the number of cells, lower the external resistance between cells, resulting in a significant decrease in the IC_50_ value. Fig 7 shows the toxicity of CdSO_4_ and 20 nm SiO_2_ nanoparticles as a function of the number of HepG2 cells grown on three different scaffolds in the absence of serum. Based on the DLS results shown above, it was confirmed that the change in the size of the nanoparticles was not large in SF media and that the cytotoxicity appeared to be greater than when serum was present due to cell uptake. In the case of 1.0 × 10^4^ cells, it was confirmed that there was no toxicity to the extent that the IC_50_ value could not be determined, as in Fig 6, where serum was present, but the cell viability (%) value was slightly reduced. In addition, as the number of cells decreased, the IC_50_ value decreased significantly in alginate and collagen. Alternately, with Matrigel, the IC_50_ values could not be obtained regardless of the number of cells. Based on these results, it was found that the toxicity results of nanoparticles differ depending on the type of scaffold used in 3D culture even under identical conditions. Other studies have reported that differences in toxicity of nanomaterials in 3D culture using Matrigel, collagen, and gelatin are caused by ECM type. This is because each ECM has its own structure and function [22].

## Conclusions

To overcome the limited ability of 2D cell culture to adequately express the *in vivo* environment, this study aimed to determine the toxicity of nanoparticles using 3D cell culture technology. Among various methods for culturing cells in 3D, a method of distributing cells mixed with ECM acting as a scaffold in a column was used in this study. The toxicity of nanoparticles in 3D cell culture method used in this study was compared to 2D cell culture. The results showed that the HepG2 cells grown in 3D are less susceptible to toxicity regardless of protein-corona formation. In addition, it was found that there were differences in toxicity according to the scaffold (ECM) type and cell number suggesting that 3D cell culture needs more research and development. Cells grown in 2D are designed to mimic *in vivo* conditions. However, the environment is significantly different with regard to morphology, exposed surface area, and intercellular signals and interactions. Previous reports have revealed that the evaluation of nanoparticle toxicity between 2D and 3D cultured cells is different, and the results of 3D cell culture are similar to *in vivo* environments and closely related to animal experiments [23]. Thus, in this study, 3D cell culture technology can also help to bridge the gap between the toxicity results of traditional *in vitro* 2D cell culture and *in vivo* assays, and is useful for developing experimental systems similar to actual *in vivo* conditions. In addition, based on the experiments conducted in this study, in future we aim to evaluate the toxicity of various nanoparticles using 3D cell cultures grown over long term to form single spheroids, which would mimic the *in vivo* environment more precisely.

## Acknowledgments

This research was supported by the Nano Material Technology Development Program [grant number 2016M3A7B6908929] of the National Research Foundation (NRF) of Korea funded by the Ministry of Science and ICT (MSIT) and Development of Measurement Standards and Technology for biomaterials and Medical Convergence funded by Korea Research Institute of Standards and Science (KRISS – 2021 – GP2021-0004).

## Supporting information

**S1 Fig.**
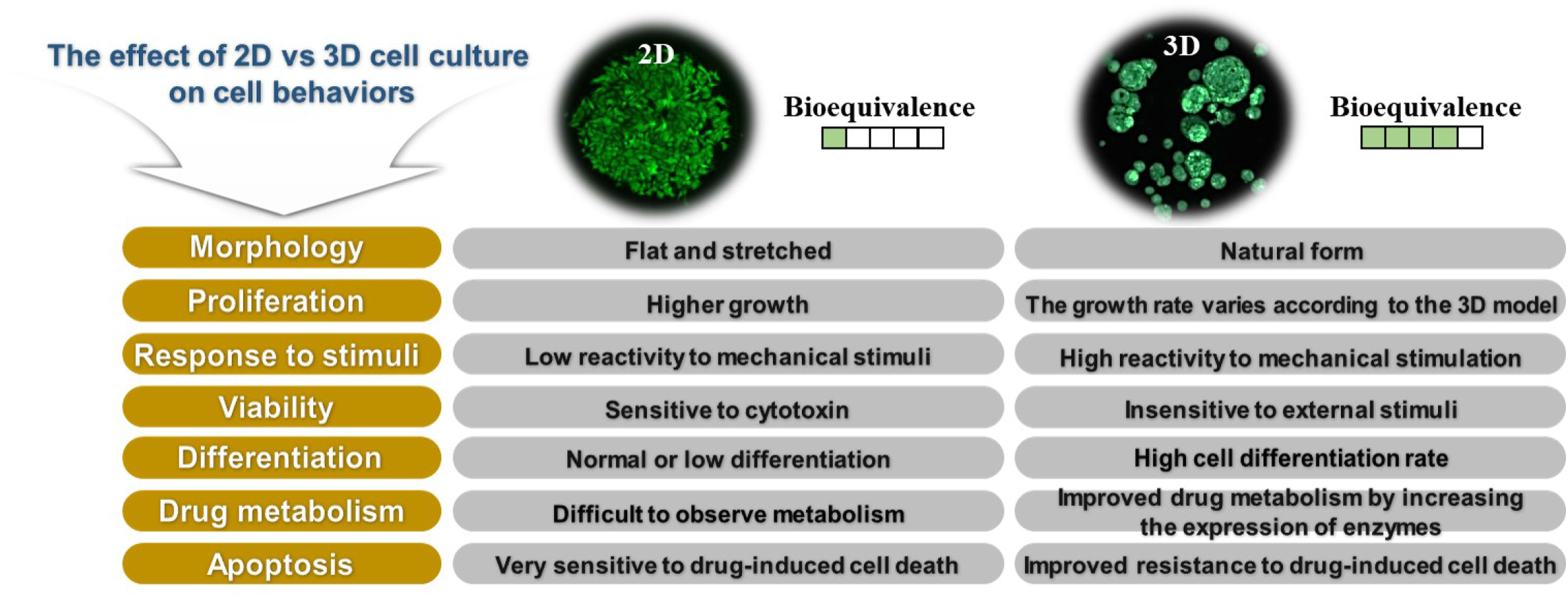
Comparison of 2D and cell culture characteristics.

**S2 Fig.**
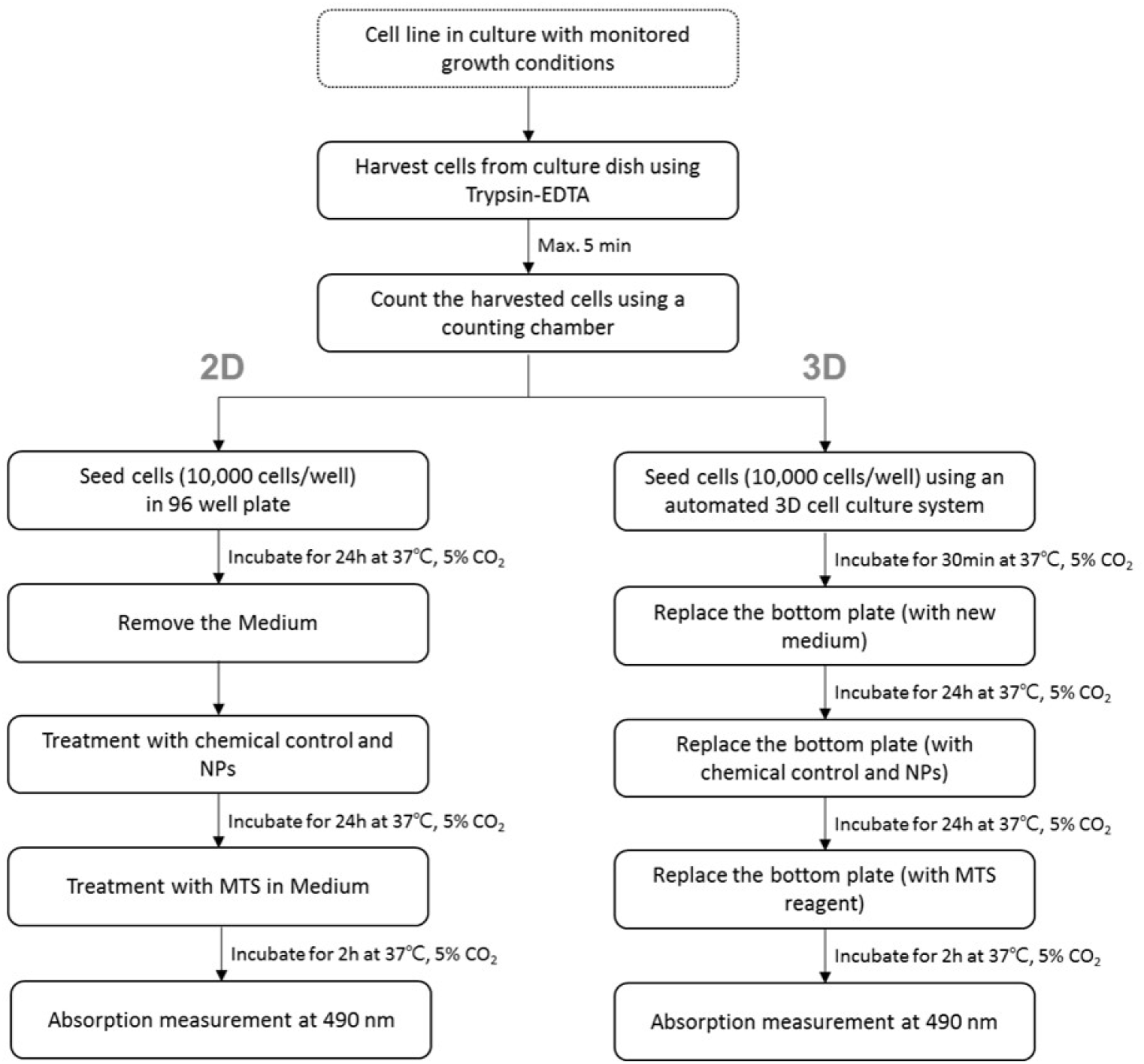
Shows the overall protocol 2D and 3D cell culture experiments.

**S3 Fig.**
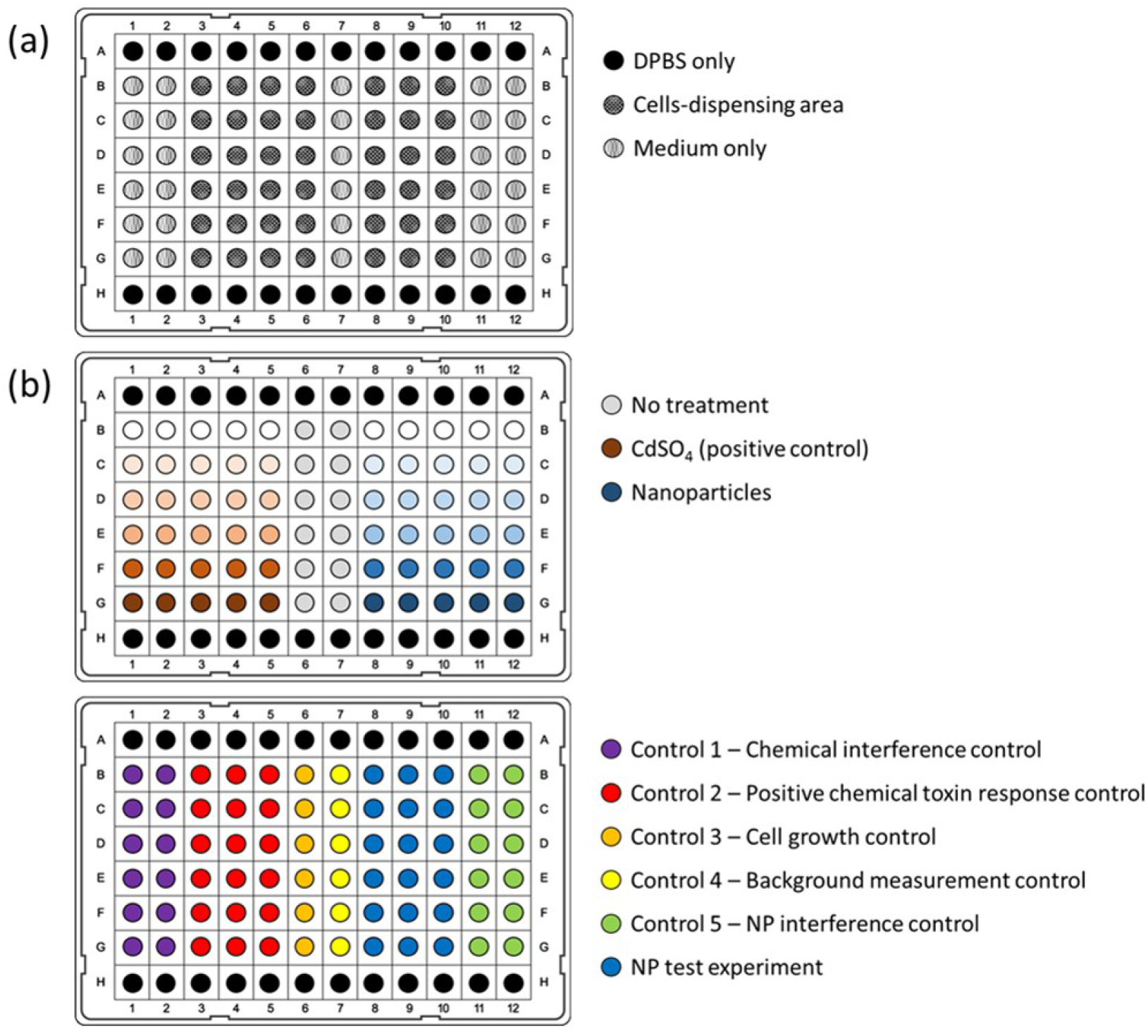
**(a)** Shows the lay out of the protocol for column/ well plates, **(b)** Shows the concentration and chemical control of the nanoparticles used in the experiment, and the control for each well is show.

